# Introgression of the sunflower *HaHB4* gene in modern wheat: An advancement in resilience to deal with climate change

**DOI:** 10.1101/2024.10.26.620406

**Authors:** Francisco Ayala, Facundo Curin, Martín Diaz-Zorita, José Murguía, Enrique Montero Bulacio, Margarita Portapila, María Elena Otegui, Raquel Lía Chan, Fernanda Gabriela González

## Abstract

Breeding for improved tolerance to water deficit is critical to mitigate climate change impact on wheat yield. In 2020, Argentina approved the first wheat transformed with the sunflower *HaHB4* gene (INDØØ412-7), which increased yield by 16% compared to the non-transformed parental Cadenza under drought conditions. Previous studies may have overestimated *HaHB4* benefits, as Cadenza is a long cycle for the Pampas region. Quantifying the yield advantage of *HaHB4* in modern, well-adapted cultivars and elucidating the physiological processes involved, is crucial for identifying optimal environments for this technology. In a broad network of experiments (one greenhouse and 29 field environments), an *HaHB4*-introgressed line of Algarrobo was compared with the conventional cultivar. Under water deficit during the reproductive phase, the yield advantage of *HaHB4* was 15% in the greenhouse and 13% in the field. *HaHB4* improved the relative yield by 0.06 to 0.08 % per mm of water deficit. The enhanced water use and water use efficiency conferred by *HaHB4*, allowed maintaining growth and yield under water deficit. *HaHB4* showed the highest benefit with moderate heat stress (∑Tmax > 30 °C ∼40-60 °Cd). In areas prone to drought combined with heat stress, *HaHB4* would enhance yield stability by improving water-limited yield.

**Highlights:** *HaHB4*-introgressed wheat enhanced water-limited yield by maintaining the crop growth under moderate to severe water deficit combined with mild heat stress through increasing water use and water use efficiency.

## Introduction

Wheat is one of the most important cereals for food security (Tweeten and Thompson, 2008; Chand, 2009). Recent climate change has already decreased wheat yield and production, (Magrin et al., 2009; Ray et al., 2019), and the climate extremes explain half of the variability of global yield anomalies (Vogel et al., 2019). Future unfavourable climate scenarios, particularly those including greater temperatures and lower and more erratic rainfall, will affect water availability for crop growth and yield generation (Hatfield and Dold, 2019). Breeding for enhanced water productivity is crucial to maintain, or even increase, yield genetic gain and production in coming years (Condon et al., 2004; Passioura, 2006; Hatfield and Dold, 2019). In 2020, the first drought tolerant transgenic wheat, the “HB4 wheat”, was approved in Argentina for food, feed and cultivation, followed by several approvals in different countries (https://www.isaaa.org/).

The “HB4” refers to the sunflower *HaHB4* gene, which encodes a divergent homeodomain-leucine zipper I transcription factor (Arce et al., 2011). It was expressed as a transgene in Arabidopsis, soybean, and wheat conferring them drought tolerance (Dezar et al., 2005; González et al., 2019; Ribichich et al., 2020). Although the molecular mechanisms involved are largely unknown, in Arabidopsis, the inhibition of ethylene perception not involving ABA-mediated stomatal closure could partially explain the differential phenotype (Manavella et al., 2006). Moreover, in wheat (IND- ØØ412-7) the heterodimerization of HD-Zip I members from different species was shown, indicating the possibility of inhibition or activation of endogenous proteins from this family or the blocking of cis-acting regulatory elements in the targets of these transcription factors (González et al., 2019).

The wheat cultivar transformed to express the *HaHB4* gene was Cadenza, generating the line INDØØ412-7 (Cadenza-HB4). This line outyielded Cadenza control by 6% average when sown in 37 field trails covering a wide range of environments within the Argentine Pampas (González et al., 2019). The mean yield gain from *HaHB4* rose to 16% when only dry environments were considered (i.e. those with a negative water balance during the reproductive period) (González et al., 2020). Similarly, when soybean Williams 82 was modified to express the *HaHB4* gene, a 4% average yield increment was observed (Ribichich et al., 2020), reaching 8.6% when only dry conditions were considered (González et al., 2020). The crop physiological processes contributing to the drought tolerance are relatively well understood in soybean but incipiently known in wheat.

Yield can be understood as the product between (i) the grain number produced per m^2^ (GN) and the individual weight of those grains (GW); or (ii) the accumulated crop growth or total biomass produced at maturity (BT) and the proportion of that biomass allocated to grains, known as the harvest index (HI). In both wheat and soybean, the mentioned field scale benefits conferred by *HaHB4* were mainly associated with improved GN and BT, without any impact on phenology (González et al., 2019; Ribichich et al., 2020). The BT depends on the interception of photosynthetic active radiation and leaf photosynthesis (Loomis and Connor, 1992). Alternatively, under water deficit, yield would depend on the amount of water used by the crop (WU, or crop evapotranspiration) and the water use efficiency (WUE) to produce biomass (WUE_BT_) and yield (WUE_YIELD_) (Passioura, 2006). In soybean, greater interception of radiation during the reproductive phase (also observed as prolonged stay-green), and enhanced leaf photosynthesis conferred by *HaHB4* contributed to increased biomass. Additionally, higher WU, WUE_BT_ and WUE_YIELD_ further supported yield increase (Ribichich et al., 2020). Noteworthy, *HaHB4* expression in soybean increased WU not only under water deficit but also under well-watered conditions. The Cadenza-HB4 showed greater efficiency to produce yield per mm of rainfall (9.4% under all environments explored and 14.2% in low-rainfall environments) (González et al., 2019), but the precise response of WU and WUE remained unknown. Given the significant differences between wheat and soybean (cereal-monocot vs legume/oil-dicot), and their distinct growing seasons (winter-spring vs spring-summer), the crop responses to *HaHB4* expression may not be analogous. Understanding the physiological processes underlying the yield advantage, particularly those related to WU and WUE, is crucial for identifying the environments where *HaHB4*-transformed wheat could provide the greatest benefits.

The genetic background in which *HaHB4* is expressed may modify the yield benefit. Because Cadenza cultivar can successfully undergo biolistic transformation (Sparks and Jones, 2004), it was selected for *HaHB4* transformation. Cadenza was released by CPB Twyford to the UK market in 1995 and has never been commercialized in Argentina. It is not well adapted to the mid latitude (28-38°) of the Pampas region because its anthesis takes place 20 to 30 days later than local commercial cultivars (Supplementary Fig. S1), exploring the reproductive phase less favourable conditions. Consequently, previous research with Cadenza-HB4 (González et al., 2019) may have helped to overexpress the benefits of *HaHB4*. Then, we wondered if *HaHB4* transformation advantages were conserved in well-adapted cultivars.

In this work we studied the impact of *HaHB4* introgession in the intermediate-late Algarrobo cultivar (Supplementary Fig. S1), which was released to the Argentine market in 2014 (https://gestion.inase.gob.ar/registroCultivares/publico/catalogo). It was highly sown by farmers during the last years, reaching ca. 1 million ha by 2020 (https://www.argentina.gob.ar/sites/default/files/sisa_if_trigo2021.pdf). A broad network of experiments, comprising one greenhouse and 29 field environments, was conducted from 2018 to 2022. The objectives were (i) to quantify the putative yield benefits of *HaHB4* introgression in a modern, well-adapted cultivar and (ii) to elucidate the physiological processes involved in this advantage.

## Materials and Methods

### Plant Material

Two cultivars were used in all the experiments: Algarrobo (AG) and an *HaHB4-* introgressed line derived from it (AGHB4). AG was crossed to IND- ØØ412-7, and the F1 was back-crossed to AG twice to obtain the back-cross 2 (BC2). In the back-cross 1, ∼480 Single Nucleotide Polymorphisms (SNP) assisted in recovering AG background. After the first self-pollination of BC2 (BC2S1), phenotypic recurrent selection for the AG parent was performed during four generations, finally obtaining the AGHB4 line (Supplementary Fig. S2)

### Experiments

One greenhouse and 29 field experiments were carried out from 2018 to 2022 (Supplementary Fig. S2). In all the experiments, pests and diseases were avoided by spraying pesticides and herbicides during the growth cycle.

The greenhouse experiment was performed in 2019 at the Rizobacter Experimental Site in Pergamino (−33.88 lat. −60.56 long), Buenos Aires, Argentina. AG and AGHB4 were evaluated under two water regimes: well-watered (WW) and water-deficit (WD). Containers of approximately 1.44 m^3^ were filled with a mix of soil/sand (4/1), except for the bottom 10 cm, which was filled with gravel. Eight rows per container, spaced 0.15 m apart, were sown at 300 plants m^-2^ on 21 June (Supplementary Fig. S3). Soil water was measured using a Diviner 2000 probe (Sentek Pty Ltd, Australia) through an access plastic tube installed in the center of each container before sowing (Supplementary Fig. S3). The available nitrogen in the uppermost 60 cm soil layer was set to 107 kg N ha^-1^ by adding urea at the beginning of tillering (24 July). All the containers were similarly irrigated to reach 85% soil field capacity until the differential water regimes started on 15 August, when the crop reached the tillering stage. Soil water measurements were performed twice weekly, each 10 cm up to 1 m depth. For the WW condition, total soil water at 1 m depth was kept at 160 mm while for the WD condition, it was set at 100 mm (Supplementary Fig. S3). Irrigations to maintain the water levels were done twice to four times a week, totalizing 653, 630, 341, and 419 mm for AG-WW, AGHB4-WW, AG-WD, and AGHB4-WD, respectively. The containers were arranged in a complete block design, with three replications distributed according to the greenhouse borders and the sun’s path. Crops were grown under the natural daylength and incoming radiation of the season. The maximum temperature in the greenhouse was set to 25°C.

The first set of field experiments aimed at studying the phenological response of the cultivars under rainfed conditions (Supplementary Table S1). AG and AGHB4 were sown in Pergamino in four (2018) or five (2019) sowing dates ranging from mid-May to the end of July, totalizing 9 environments (Supplementary Table S1). The plant population was adjusted to the sowing date ranging from 200 (earliest) to 360 plants m^-2^ (latest). The second group comprised yield comparative experiments in different locations across the central region of the wheat production area (−32.4 to −38.4 latitude, and −58.2 to −64.3 longitude, Supplementary Fig. S2), totalizing 20 environments (Supplementary Table S1). While some experiments included drip irrigation (IR), most of them were conducted under rainfed (RF) conditions. Sowing dates ranged from 10 June to 2 July for locations in the northern and west area (Arias, Monte Buey, Pergamino, Roldan, Victoria, Santa Rosa) and from 17 June to 5 August for locations in the southern area (Balcarce, Tandil, Tres Arroyos, Bordenave) (Supplementary Table S1, Supplementary Fig. S2). The plant population was always 240 plants m^-2^. In the northern and west area, the fertilization was 100 N + 40 P + 30 S + 1,2 Zn kg ha^-1^ at sowing, with an additional 200 Kg N ha^-1^ at tillering. A similar fertilization rate was used in the southern area, except for the S and Zn that were not added.

All field experiments were machine-sown in 7-row plots, 0.20 m apart and 6 m long. Plots were arranged in a complete block design with three replications. When irrigation was present (drip method), two independent sites were established (irrigation -IR and rainfed- RF), and the blocks were nested within each site. A similar arrangement was performed for each date within the sowing date experiment. Each combination of location × year × treatment (rainfed/irrigation/sowing date) was considered one environment (Supplementary Table S1).

### Measurements

Detailed phenology was measured particularly in the greenhouse and sowing date experiments in Pergamino. The occurrence of emergence (Z1.0), first node detectable (Z3.1), flag leaf expanded (Z3.9), beginning of anthesis (Z6.1), and physiological maturity or yellow peduncle (Z8.7) were measured following Zadoks scale (Zadoks et al., 1974). These stages, except for flag leaf expanded, were also measured in the rest of the experiments performed in Pergamino as key developmental stages to take other samples (see below). The date of first node detectable was set when 50 % of 20 random plants had the first detectable node, while the rest of the stages were considered when ca. 50% of the plants in a plot reached that stage (general visual assessment).

To estimate yield, and its numerical components and physiological determinants, different samplings were performed according to the type of experiment. For the field experiments conducted in Pergamino and Santa Rosa, when each plot reached maturity (Z9.2), plants in two meters long central rows were cut at their base to estimate the harvest index (HI) and the individual grain weight (GW). The spikes were counted to estimate their number per m^2^ and separated from the rest of the plant. Both fractions were dried in an air-forced oven at 70°C until constant weight. Once the fractions were weighed (stover and spikes with grains), they were summed up to calculate the total biomass of the sample. The spikes were hand-threshed, and the grains were weighed to calculate the sample yield. The individual grain weight (GW) was estimated as the quotient of a 20 g sample of grains divided by the grain number present in it, which were counted with an automatic machine (Pfeuffer GmbH Seed Counter). The HI was estimated as the quotient between the sample yield and the sample total biomass. The yield per m^-2^ and % humidity were obtained from the hand-harvest of all the plants present in each container or in the 2 m long rows sample in the sowing date experiments, following stationery machine-threshing. In the rest of the field experiments, it was estimated from the machine harvest. The final yield was always expressed at 14% humidity. The GW was corrected to 14 % humidity to obtain the grain number m^-2^ (GN) as the quotient between yield and GW. For Balcarce, Bordenave, and Tandil locations, as well as one of the experiments carried out in Pergamino in 2020 (PE20), the GW was calculated as described above but the subsamples of 20 g each were taken from the bulk of grains in each machine-harvested plot. The total biomass produced per m^2^ at maturity (BT 14% humidity) was calculated as the quotient between plot yield and estimated HI, and the grain number per spike as the quotient between GN and spike number per m^2^. The main determinants of this last trait, i.e. the maximum tiller number and the tiller survival score, were estimated in the field experiments performed in Pergamino during 2018, 2019, and 2020. The maximum tiller number was counted when plots reached first node detectable in a half meter length of a central row. The tiller survival was calculated as the quotient between spike number per m^2^ and maximum tiller number and expressed in percentage.

When the field experiments performed in Pergamino (except for PE20) reached anthesis, the biomass per m^2^, the spike dry weight per m^2^, and the biomass partitioning to the spike were calculated. For this purpose, plants in 1 m long central row were cut at their base and the spikes were separated from the rest of the biomass. Both fractions were dried in an air-forced oven at 70°C for 72 h and weighed to estimate the spike and biomass dry weights per m^2^ (i.e., the last one as the sum of the spikes and the rest of the biomass). The biomass partitioning to the spike was calculated as the quotiente between the spike and biomass dry weights. The fruiting efficiency was computed as the quotient between GN and spike dry weight per m^2^.

Water use (WU, or crop evapotranspiration) and water use efficiency (WUE) were estimated for the containers in the greenhouse experiment as well as the field experiments in Pergamino (except for PE20). In the containers, the water content of the soil was measured using a Diviner 2000 probe, as previously explained. In the field experiments, it was measured gravimetrically, taking samples each 30 cm up to 1.5 m (2018, 2019, 2020) or 2 m (2022) soil depth using a soil auger. The soil samples were dried in an air-forced oven at 100°C and the gravimetric water content was calculated as the difference between the humid and dry weights of the sample, relative to its dry weight. The samples were taken at emergence, first node detectable and at the end of the growth cycle (i.e. 10 days after yellow peduncle). The variation in soil water between these stages (Δ soil water) and the sum of rain and irrigation during each period were considered for the estimation of WU. The WUE_BT_ and WUE_YIELD_ were estimated as the quotient between biomass or yield to WU over the entire growth cycle, from emergence to maturity. The Normalized Vegetation Index (NDVI) was measured along the crop cycle to estimate canopy cover and greenness in the experiments performed in Pergamino during 2022, where rainfed (PE22RF) and two irrigation levels were set (PE22IR1 and PE22IR2, 220 and 548 mm, respectively). The canopy temperature of all plots was measured close to anthesis (12 and 27 Oct) and at the beginning of grain filling (4 and 8 Nov) using a HikMicro thermal camera (Serie M10, https://www.hikmicrotech.com) during 2022. To estimate canopy temperature without the influence of the soil, the pixels of the five central rows of the plot were manually selected to avoid the soil between rows. Thermal images were analysed with the Hikmicro Analyzer (2021).

In the greenhouse, some other detailed measurements were performed. The leaf area index at anthesis was estimated from a 20 cm central row sample using the LI-3000C area meter (LI-COR). The maximum leaf photosynthesis of the two uppermost leaves (flag leaf and flag leaf-1) was measured at anthesis and 15 days post-anthesis using an LI-6400/XT Photosynthesis System (LI-COR). The stomatal conductance at flag leaf emergence was measured in the same two uppermost leaves using an LI-600 porometer (LI-COR). To investigate whether AG and AGHB4 cultivars can be discriminated under the different water treatments by proximally-sensed canopy hyperspectral data, a compact shortwave near-infrared (NIR) spectrometer (STS-NIR, Ocean Insight) was used. This spectrometer can detect 1024 wavelengths within the range of 632 to 1125 nm. All measurements were performed around solar noon, between 10:00 and 14:00 ART (UTC 03:00), with the spectrometer positioned directly above the canopy surface at a distance of 50 cm, resulting in a footprint diameter of approximately 26 cm. Before each canopy measurement, the upwelling light reflected from a white reference material measuring 40 cm × 40 cm with a reflectance of 99 % was recorded. Dark current was determined by occluding the spectrometer’s entrance slot and subsequently subtracted from both white reference material and canopy measurements. Integration time was adjusted to prevent saturation of the white reference material signal (Arias et al., 2021). Ten measurements were taken per container with an even distribution, avoiding edge measurements to prevent boundary errors. Data were collected on 20 September (18 days before anthesis), 8 October (during anthesis), and 7 November (during grain filling).

The rains were recorded in weather stations placed close to experiments (less than 50 m away), whereas the air temperature and the photosynthetic active radiation (PAR) were downloaded from the NASA Prediction of Worldwide Energy Resources (https://power.larc.nasa.gov).

### Analyses

In the greenhouse containers, the ANOVA was used to test the impact of the water regimes (WW - WD) and cultivars (AG and AGHB4) on traits measured according to the following model (Eq.1):

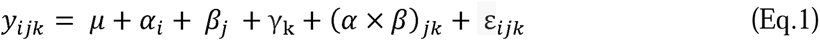

Where μ is the grand mean; *α*_i_ is the effect of the ith block (i = 3); *β*_j_ is the effect of the jth irrigation treatment (j = 2), *γ*_k_ is the effect of the kth cultivar (k = 2). The term between parenthesis (*α* × *γ*)*_jk_* corresponds to the treatment by cultivar interaction and ε_ijk_ is the error.

For the field experiments, the model used for ANOVA analyses considered the impact of environments (combination of location × year × treatment) and cultivars (AG and AGHB4) according to equation 2:

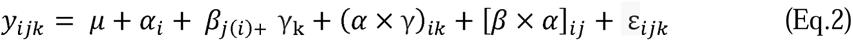

Where μ is the grand mean; *α*_i_ is the effect of the ith environment (i = 29); *β*_j(i)_ is the effect of the jth block nested within the environment (j = 3), *γ*_k_ is the effect of the kth cultivar (k = 2). The term (*α* × *γ*)*_ik_* corresponds to the environment by cultivar interaction (E×C); [*β × γ*]*_ij_* is the error (a); ε_ijk_ is the error (b).

The relative yield (RY) of AGHB4 compared to AG was calculated as

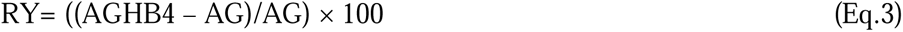

In the case of the field experiments, three ANOVAs were performed. The first one with the complete set of environments (27 environments, as 2 out of 29 were damaged by hail after anthesis), the second one with the environments where RY>0 and the third one with those where the RY<0. This partitioned analysis eliminated the interaction with the environments, helping to understand the physiological processes behind the different yield responses.

The water deficit of the environment (WDE) was calculated as the subtraction of reference evapotranspiration (ET_0_, Hargreaves and Allen, 2003) from the rain plus irrigation during the months when the reproductive phase takes place. These months are September, October and November (SON) for the northern and west locations (Pergamino, Arias, Monte Buey, Roldan, Victoria, Santa Rosa) and October, November and December (OND) for the southern locations (Balcarce, Bordenave, Tandil, Tres Arroyos). For the experiments where plant available soil water was measured at first node detectable, the water deficit during the reproductive period was calculated by subtracting the accumulated ET_0_ from the total of rainfall plus irrigation, and then adding the plant available soil water at the start of the reproductive period.

Sixteen vegetation indices were selected from existing literature to be calculated with the hyperspectral data. These indices were chosen based on their specific applications, including their sensitivity to crop biomass, chlorophyll status, and water stress (Supplementary Table S2). The vegetation indices including wavelengths below 880 nm such as NDVI, ND800680, NDRE, SR800735, CCCI, DSWI-4 and GLI, are associated with higher nitrogen, chlorophyll contents, canopy cover and yield as the value of the index increase (Apan et al., 2003; Tilling et al., 2007; El-Shikha et al., 2008; Bhandari et al., 2020; Raya Sereno et al., 2024). Conversely, lower values of FR and FR2 indicate better crop conditions associated with higher chlorophyll content and a high chlorophyll/carotenoid ratio (Stober and Lichtenthaler, 1992). On the other hand, the vegetation indices that consider wavelengths above 880 nm, focus on water absorption bands such as WI, PWI, WBI, WBI3, SR980900 and RATIO975. Lower values of these indices correspond to better crop conditions and have been correlated with yield (Gutiérrez et al., 2010) and canopy water content in the case of RATIO975 (Pu et al., 2003). The exception of this group is the WBI4 index, which shows an inverse relationship due to its mathematical definition. The values of vegetation indices were normalized to ensure uniform scaling. Additionally, new indices were proposed to enhance the analysis. The spectral band ranging from 680 nm to 755 nm was examined using its first-order derivative. This range was segmented into three parts (680-721 nm, 721-740 nm, and 740-755 nm) to assess the rate of change in reflectance within these specific wavelength segments. The derivatives were calculated by determining the slope between consecutive points, followed by computing the mean values for each interval (Rigalli et al., 2018), resulting in dA, dB, and dC, respectively. From these values, the ratio between dC and dB (dC/dB) was calculated, as well as the normalized differences between dA and dC (NDdA/dC) and between dB and dC (NDdB/dC). Finally, the maximum derivative value between 723 nm and 728 nm was also computed. The formulas used for these calculations are provided in the Supplementary Table S2.

The ANOVAs, the linear regressions, and Pearson correlations between measured traits were performed using InfoStat (Di Rienzo et al., 2020).

## Results

### Greenhouse experiment

#### AGHB4 outyielded AG when water was restricted under controlled conditions

The yield ranged from 500 to 673 gm^-2^ and depended on the cultivar × water regime interaction (Fig. 1A and Supplementary Table S3). Under WW, both cultivars yielded similarly, though there was a trend for a 7% higher yield in AG. However, when water was restricted, AGHB4 yielded 15% more than AG (Fig. 1A). AGHB4 showed greater yield stability, reducing yield only by 7% compared to the 26% reduction of AG under WD.

**Fig. 1.**
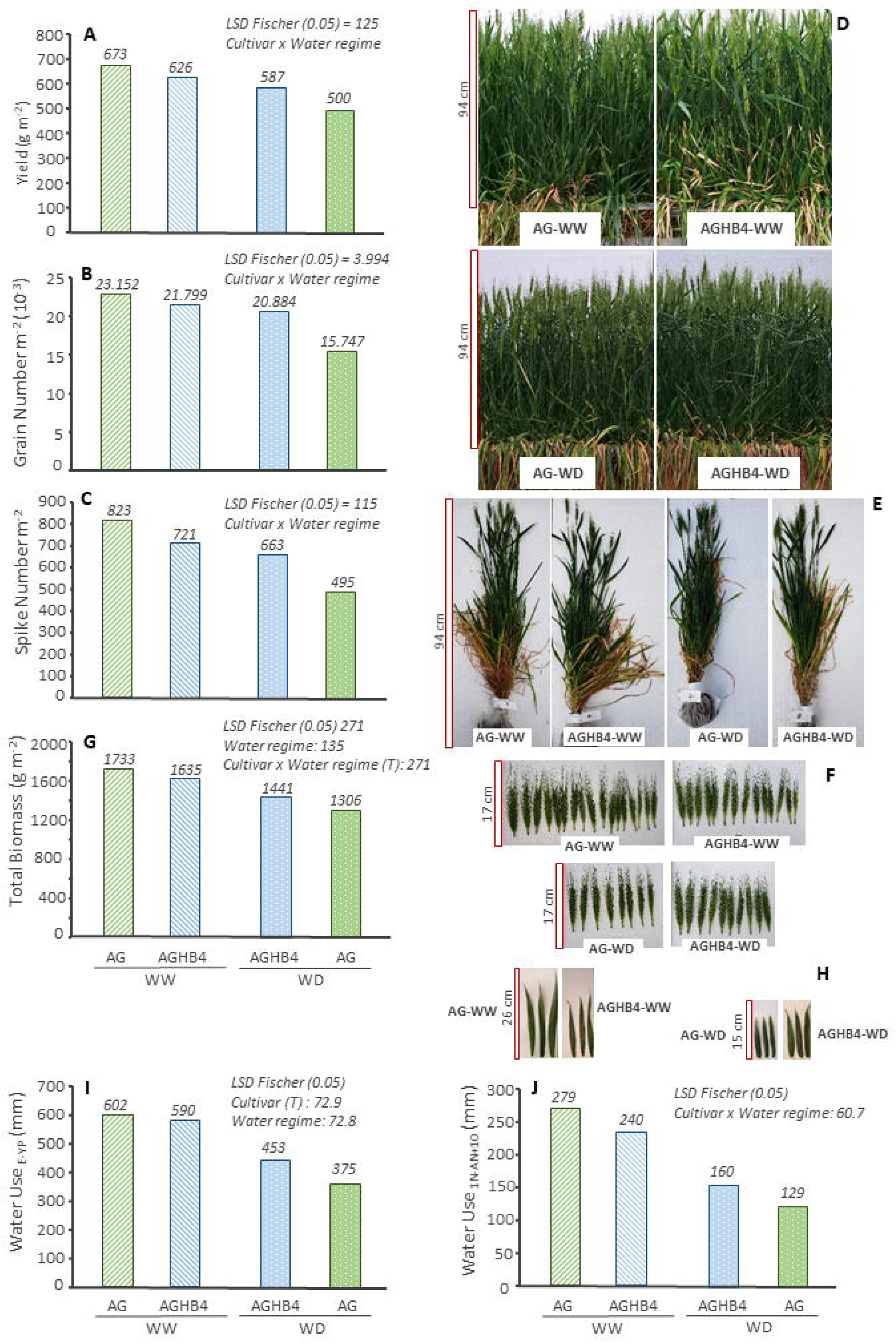
Response of Algarrobo (AG) and its *HaHB4*-introgressed line (AGHB4) to well-watered (WW) and water-deficit (WD) treatments in the greenhouse. (A) Yield, (B) grain number per m^2^, (C) spike number per m^2^, illustrative pictures of (D) plants close to anthesis, (E) plants sampled at anthesis and (F) spikes present in the sample, (G) total biomass at maturity, (H) illustrative pictures of flag leaves, (I) water use from emergence (E) to physiological maturity (or yellow peduncle-YP), (J) water use during the reproductive phase from the first node detectable (1N) to ten days post anthesis (AN+10). Note the scale (cm) in the left side of the pictures. The numbers close to each bar indicate the actual value of the variable. The LSD Fischer value (α=0.05) for the statistically significant effects is shown in the upper right corner of each sub-figure. The T indicates an statistical trend (0.05>P<0.1).

The cycle to maturity was modified by the cultivar and the water regime (p<0.05, data not shown). AGHB4 was 2 days shorter than AG (152 vs 154 days), while the WD reduced 4 days the time to maturity respect to WW (151 vs 155 days). The time from sowing to first node detectable (Z3.1) showed no difference at all (∼ 81 days), whereas the time from first node to anthesis (Z6.1) depended on cultivar and water regime (p<0.05). AGHB4 was 3 days shorter than AG (33 vs 36 days) while the WD also reduced by 3 days this phase (33 vs 36 days). For the phase from anthesis to maturity (Z8.7) no statistical differences were observed (∼36 to 39 days). These differences in duration from first node to anthesis were not significantly correlated with yield (p=0.23844).

#### The number of grains and spikes per unit area explained the differences in yield between AG and AGHB4

The yield showed a strong correlation with the GN (r=98%, p=0.0209) and no relationship with the GW (p=0.2266). Similar to yield response, GN depended on the cultivar × water regime interaction (Fig. 1B and Supplementary Table S3). When the crop was WW, AG tended to produce more GN than AGHB4 (6%) but when water was restricted, AGHB4 maintained the GN (−4%) while AG lowered it (−32%, Fig. 1B). As a result, AGHB4 produced 25% more GN than AG under WD. The GW was affected by the cultivar and tended to respond to the cultivar × water regime interaction (Supplementary Table S3). The GW was remarkably similar among treatments (∼28.3 - 29.1 mg), except for AG under WD, where it increased to 31.9 mg, likely as a trade-off to the reduced GN (Supplementary Table S3).

The GN variation was mostly associated with the number of spikes per m^2^ (r=0.98, p=0.0222), while it did not respond (p=0.3914) to the number of grains per spike. Similar to GN, the number of spikes per m^2^ depended on cultivar × water regime interaction (Fig. 1C, Supplementary Table S3). Under WW, AG tended to produce 12% more spikes than AGHB4, whereas, under WD, AGHB4 had 25% more spikes than AG (Fig. 1C and pictures in Fig. 1D, E and F). Then, when comparing WW vs WD, AG reduced 40% the number of spikes per m^2^ while AGHB4 did it only by 8%. The number of grains per spike ranged from 28.6 to 32.0 and did not differ between cultivars or water regimes (Supplementary Table S3).

#### Biomass at maturity and leaf area index were responsible of the greater yield of AGHB4 under water deficit

The yield was primarily correlated with BT (r=0.97, p=0.0267), while HI had no significant impact (p=0.8268). The BT was affected by the water regime and showed a trend to respond to the cultivar × water regime interaction resembling the responses of yield, GN and spike number per m^2^ (Fig. 1E, Supplementary Table S3). AG tended to produce 6% more BT than AGHB4 under WW, however, under water deficit AGHB4 produced 10% more BT than AG (Fig. 1G). Meanwhile, the HI tended to respond to the cultivar x water regime interaction in a different pattern, being similar between cultivars under WW, but increasing in AGHB4 under WD (0.41 vs 0.38 in AGHB4 vs AG, Supplementary Table S3).

The GN would depend on the growth of the spikes at anthesis (spike dry weight at anthesis per m^2^) and the efficiency of those spikes to set grains (or fruiting efficiency, grains per gram of spike weight). In the present assay, the GN was correlated with both traits (r=0.98, p=0.0127 for the first one and r=0.95, p=0.0437 for the second one). Although no statistical difference was detected, the response pattern of both traits was similar to that of GN (Supplementary Table S3). The leaf area index at anthesis, which reflects the crop’s ability to intercept radiation, responded to the cultivar × water regime interaction, following the same response pattern to BT (Supplementary Table S3). Under WW, AG showed 33% higher leaf area index than AGHB4, but the relationship reversed under WD, where AGHB4 showed 10% more leaf area (Supplementary Table S3). This response was also evident in the flag leaves (Fig. 1H). The photosynthesis of the upper leaves responded to the water regimes, decreasing ∼31-46% under WD (Supplementary Table S3). AGHB4 showed a trend to greater (8-9%) photosynthesis than AG under WD, but the difference was not statistically significant. Leaf stomatal conductance was not statically modified by any treatment, but the response pattern was similar to the other traits. Under WW, it seemed to be 8% greater in AG than AGHB4 and it tended to reverse under WD, showing a tendency to be 12% greater in AGHB4 than AG (Supplementary Table S3).

#### AGHB4 used more water than AG under water deficit

The WU from emergence to maturity (WU_E-YP_) ranged from *ca*. 375 to 602 mm. It depended on the water regime and tended to change with the cultivar (Fig. 1I, Supplementary Table S3). Under WW, the WU_E-YP_ was close to 600 mm, while it was reduced by 40 % when water was restricted in the WD. Averaging across water regimes, AG consumed 489 mm while AGHB4 reached 521 mm. Nevertheless, this result was highly influenced by the WU_E-YP_ of AGHB4 under WD. Despite no statistical difference being established, the WU_E-YP_ of AGHB4 under WD was 17% greater than that of AG, whereas under WW it tended to use 2% less water than AG (Fig. 1I).

When the water use was computed for the entire reproductive period (from first node detectable to maturity, WU_1N-YP_), or specifically during the grain number determination phase (from first node detectable to 10 days after anthesis, WU_1N-An+10_), the interaction between water regimes and cultivars was statistically more significant (Fig. 1J, Supplementary Table S3). Under WW, the WU_1N-An+10_ of AGHB4 was 14% lower, while under WD it was 19% greater than that of AG (Fig. 1J). This response pattern mimicked that of yield, GN, and spike number per m^2^.

The WUE_BT_ ranged from 2.8 to 3.5 g m^-2^ mm^-1^ and it was affected only by the water regime, being 14% higher under WD compared with WW (Supplementary Table S3). A similar response was observed for the WUE_YIELD_, which ranged from 1.1 to 1.3 g m^-2^ mm^-1^ (Supplementary Table S3).

#### Several vegetation indices discriminated AG from AGHB4 under water deficit

The vegetation indices related to water stress were sensitive at all three studied time points (pre-anthesis, anthesis, and grain filling) to discriminate cultivars under WD (Fig. 2). The indices related to chlorophyll status were useful for identifying differences in the pre-anthesis stage (GLI) and during anthesis (NDRE and CCCI), while the indices related to canopy cover and total biomass (NDVI and ND800680) identified differences during the grain filling. Interestingly, the vegetation indexes did not show significant differences between cultivars under the WW treatment except for WBI3 in pre-anthesis and WBI4 in anthesis (Fig. 2). The newly proposed indices, dCdB, NDdAdC, and NDdBdC, proved to be useful for identifying differences between cultivars under WD during anthesis, whereas MdD was effective during grain filling.

**Fig. 2.**
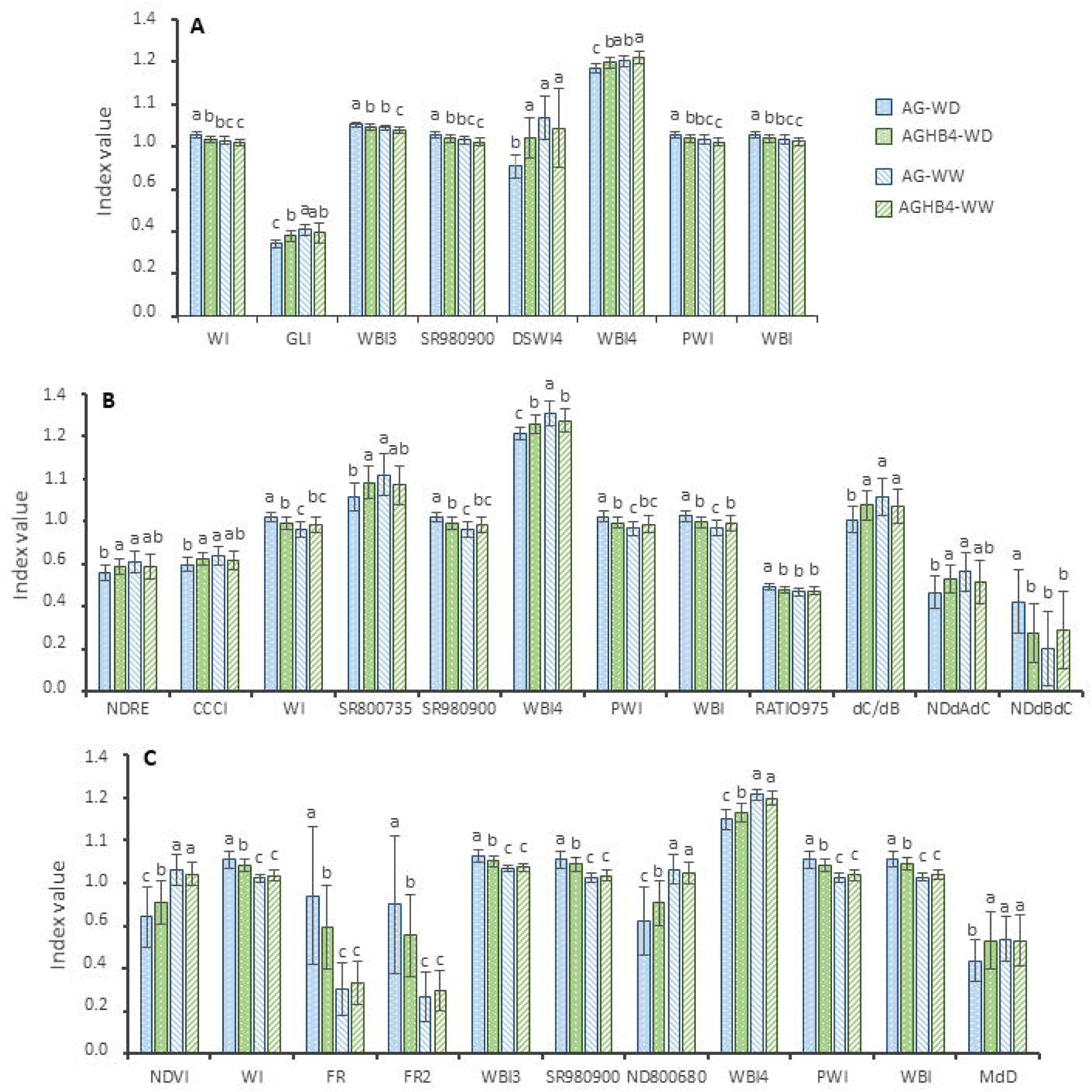
Vegetation indices discrimination. Index values (A) during pre-anthesis (−18 days), (B) at anthesis, and (C) during grain filling for Algarrobo (AG) and its *HaHB4*- introgressed line (AGHB4) under well-watered (WW) and water-deficit (WD) treatments. Different letters above error bars denote statistically significant differences (Tukey α = 0.05) among cultivars and water treatments. For the vegetation indices description see Supplementary Table 2.

### Field Experiments

The field environments explored were highly diverse. The rainfall and irrigation accumulated during the months when the reproductive phase typically occurs (SON or OND, depending on the location) ranged from 95 to 508 mm, while the accumulated ET_0_ varied from 360 to 498 mm (Supplementary Table S1). In Pergamino, the 2018 season was particularly rainy and cold, while the 2022 season was extremely dry and warm. The sum of mean daily temperatures >20 °C and maximum temperatures >30 °C were 3 and 13 times higher in 2022 than in 2018, respectively (Supplementary Table S1).

#### Similar phenology was exhibited by AG and AGHB4 in different sowing dates under field conditions

When both cultivars were tested on different sowing dates for two years, the life cycle was very similar between them. AGHB4 was slightly earlier than AG, anticipating anthesis by 2 to 6 days, depending on year and sowing date. This difference was also observed in previous stages (flag leaf 2 to 4 days, and first node 0 to 4 days) as well as in later ones (yellow peduncle 2 to 4 days, Supplementary Fig. S4). These results agree with the similar phenology described previously for the greenhouse containers and for the rest of the field experiments where anthesis was measured (Supplementary Table S1).

#### AGHB4 outyielded AG in 23 out of 27 field environments being the water deficit the driving force of such a response

Considering all field experiments, yield ranged from 1490 (SR22RF AG) to 11229 kg ha^-1^ (PE18SD1 AG) depending on the environment, cultivar, and the cultivar × environment interaction (p<0.05, Fig. 3A, Supplementary Table S4). In only 4 out of 27 environments, AG yielded higher than AGHB4. This occurred in the highest yielding environments (PE18SD1/PE18SD2/PE18SD3) or when rain plus irrigation was among the highest records (PE22IR2, 508 mm accumulated during SON). Averaging all environments, the yield of AGHB4 was 8.1% higher than that of AG (5215 vs 4791 kg ha^-1^, Supplementary Table S4). Nevertheless, this general average concealed a wide range of relative responses between both cultivars, depending on the environment. The relative yield (RY) of AGHB4 compared to AG ranged from −12.0 to +37.3% (Fig. 3B, Supplementary Table S4). No relationship was observed between the RY and the differences in days to anthesis between AGHB4 and AG (r=0.35, p=0.2618). The driving force for the RY response was the water deficit of the environment (WDE) during the months when the reproductive phase took place. The WDE explained 27% of the variation in the RY, showing AGHB4 a 0.06 % increase in RY per mm of WDE (p=0.0034) (Fig. 3B). Then, for the environments with RY>0 (all of them with WDE<0 except for SR22IR) the average yield advantage of AGHB4 was 13% (4784 vs 4164 Kg ha^-1^, p<0.00001, Supplementary Table S4). Meanwhile, for the rest of the locations with RY<0 the differences in yield were not statistically significant (9.1% RY, 7689 vs 8393 kg ha^-1^, AGHB4 vs AG, Supplementary Table S4).

**Fig. 3.**
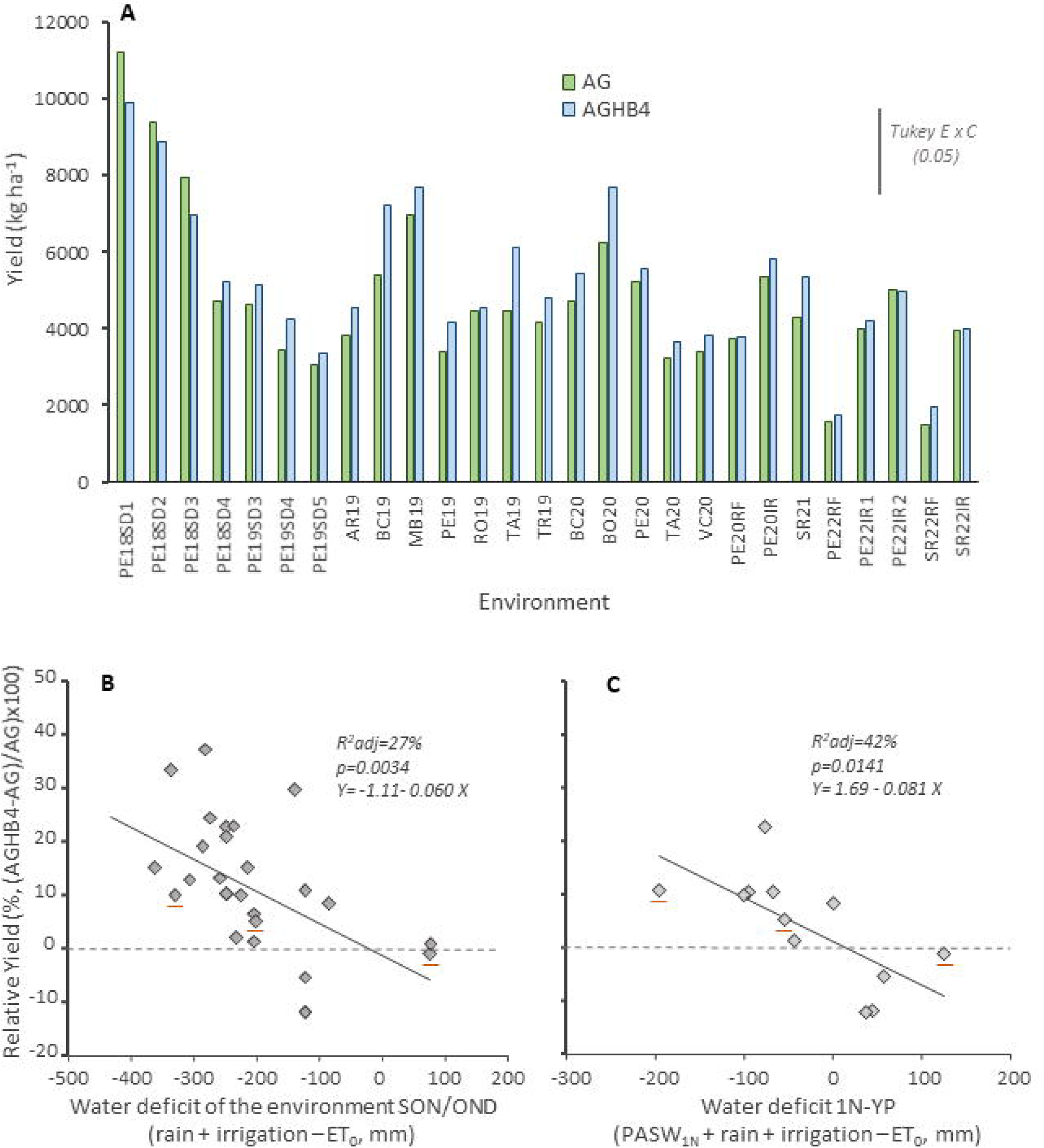
Yield response under field conditions. (A) Yield for each environment and cultivar. The vertical line in the upper right corner represents the Tukey value (α=0.05) for the environment × cultivar interaction. (B) Relative yield of AGHB4 over AG related to the water deficit of the environments during the months when the reproductive phase takes place (September, October and November for northern locations -SON, and October, November and December for the southern locations - OND). The water deficit was estimated as the rain + irrigation - evaporative demand (ET_0_). (C) Relative yield of AGHB4 over AG related to the water deficit during the reproductive phase from the first node detectable (1N) to maturity (or yellow peduncle- YP). Water deficit was considered as the balance between the plant soil available water up to 1.5 or 2 m at 1N (PSAW_1N_) + rain + irrigation – ET_0_. This water balance was calculated for AG as the control cultivar. The three underlined dots in (B) and (C) represent the response for PE2022 rainfed and the two irrigation levels. AG: Algarrobo. AGHB4: *HaHB4*-introgressed line derived from Algarrobo.

For Pergamino, the water deficit during the reproductive phase (from first node to yellow peduncle) was calculated considering the plant available soil water at first node detectable. In this case, the yield advantage of AGHB4 per mm of deficit increased to 0.081% (p=0.0141, R^2^=42%) (Fig. 3C). Interestingly, in the four previously mentioned environments where AG outperformed AGHB4, there was no water deficit when considering the plant available soil water. This reinforced the idea that water deficit is the key factor behind the yield advantage of AGHB4. Only in one environment (PE20IR) with a positive water balance (0.76 mm) during the reproductive phase, a positive yield response (8.5%) of AGHB4 compared to AG was computed (Fig. 3C).

#### The yield benefit of AGHB4 was explained by the grain and spike number per m^2^

Considering all environments and both cultivars, the GN ranged from 4946 to 29812 grains m^-2^ (AG in PE22RF and PE18SD1, respectively, Supplementary Table S5). In turn, the GW varied from 19.9 to 39.0 mg (AGHB4 in PE19SD5 and AG in PE18SD2, respectively, Supplementary Table S5). The yield variation was mostly explained by the GN (R^2^= 78%, p<0.0001, n=19), while a reduced variation was associated with the GW (R^2^=26 %, p=0.0006, n=19).

When RY>0, GN and GW depended on the environment and cultivar, with no interaction effect observed (Supplementary Table S5). Averaging all environments with RY>0, AGHB4 showed 7.1% more grains and 3.7% heavier grains than AG (Fig. 4A and B). No statistical difference was detected in GN and GW between cultivars for the environments with RY<0 (22504 vs 23920 grains m^-2^ and 33.7 vs 34.8 mg for AGHB4 vs AG, Supplementary Table S5).

**Fig. 4.**
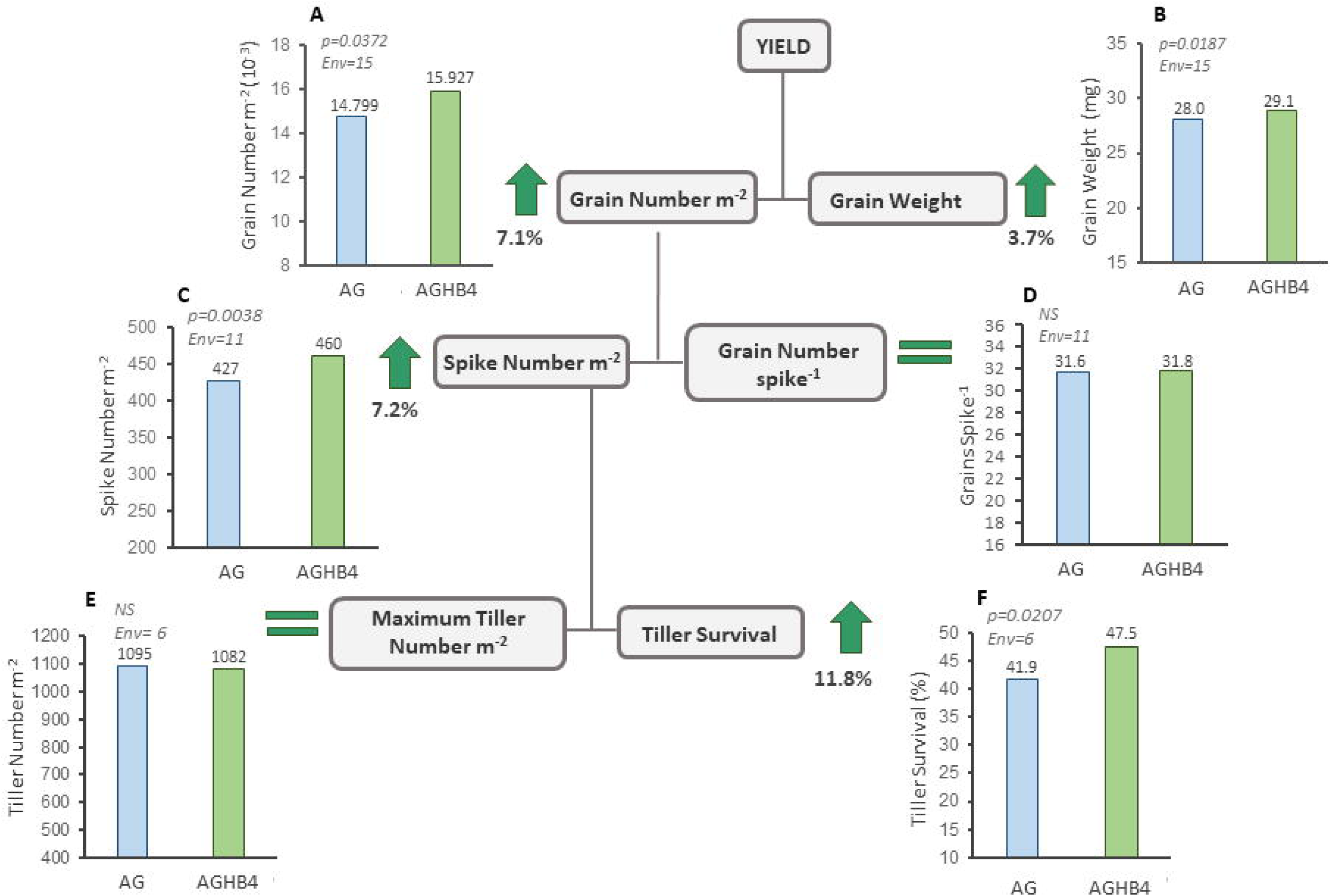
Numerical components of yield in the environments where the relative yield (RY) of AGHB4 over AG was >0. (A) Grain number per m^2^, (B) individual grain weight, (C) spike number per m^2^, (D) grain number per spike, (E) maximum tiller number per m^2^, and (F) tiller survival. The numbers above the bars indicate the exact value of the variable. In the upper left corner of each sub-figure, the p-value and the number of environments (Env) in which the trait was tested are displayed. AG: Algarrobo. AGHB4: *HaHB4*-introgressed line derived from Algarrobo.

Variation in GN was associated with both the spike number per m^2^ (R^2^=62 %, p<0.0001) and the grain number per spike (R^2^=77 %, p<0.0001) when all environments and both cultivars were considered together. The spike number per m^2^ ranged from 264 to 641 (AGHB4 in PE22RF and PE18SD2, respectively, Supplementary Table S5). It was affected by the environment and cultivar when RY>0 (Supplementary Table S5). In this case, AGHB4 set an average of 7.2% more spikes than AG (Fig. 4C). The relative number of spikes per m^2^ of AGHB4 compared with AG (calculated similarly to RY), increased by 0.04% per mm of WDE (R^2^=33%, p=0.0491, n= 10, all environments RY>0, except for PE22RF where there was an extreme low spike number per m^2^, Supplementary Table S5). The grain number per spike varied from 17.1 to 53.9 (AG in PE22RF and PE18SD1, respectively) and was solely modified by the environment, with no impact of cultivar (Fig. 4D, Supplementary Table S5).

The spike number per m^2^ is the result of the maximum number of tillers developed and the survival of those tillers to produce a spike. The maximum number of tillers was only modified by the environment (ranging from 629 to 1715 tillers m^-2^, AG in PE20RF and AGHB4 in PE18SD2, respectively) with no cultivar impact (Fig. 4E). Meanwhile, the survival of those tillers was affected by the environment and the cultivar, particularly when RY>0 (Supplementary Table S5). In these cases, AGHB4 showed a tiller survival 11.8% greater than AG (Fig. 4F). In contrast, when the RY<0, no difference was observed between cultivars or among environments for tiller survival (Supplementary Table S5).

#### The biomass at maturity and the spike dry weight at anthesis support the increased grain number and yield of AGHB4

The BT ranged from 3077 to 22319 kg ha^-1^ (AG in PE22RF and PE18SD1, respectively, Supplementary Table S6), whereas the HI ranged from 0.29 to 0.55 (AG in SR22RF and AGHB4 in PE20IR, Supplementary Table S6). Considering all data, yield variation was mostly explained by the BT (R^2^=89%, p<0.0001) with a reduced impact of HI (R^2^=10%, p=0.0432). For environments where RY>0, both the HI and BT depended on environment and cultivar (Supplementary Table S6), showing AGHB4 higher BT (6.9%) and HI (2.3%) than AG (Fig. 5A and B). No statistical differences were observed between cultivars for these traits when RY<0, though a significant effect of the environment was computed (Supplementary Table S6).

**Fig. 5.**
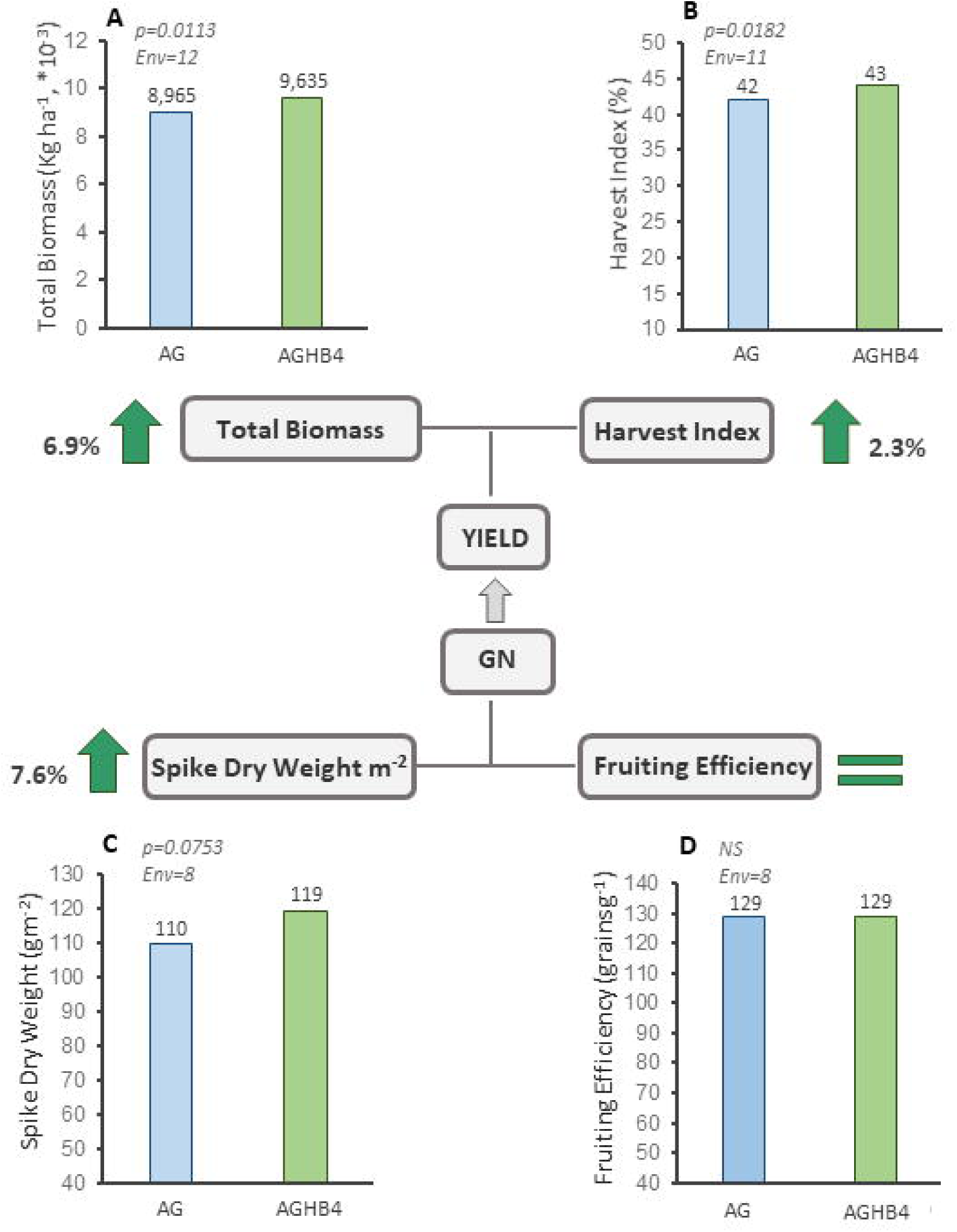
Crop physiological determinants of yield in the environments where the relative yield (RY) of AGHB4 over AG was >0. (A) Total biomass at maturity, (B) harvest index, (C) spike dry weight at anthesis, (D) fruiting efficiency. The numbers above the bars indicate the exact value of the variable. In the upper left corner of each sub-figure, the p-value and the number of environments (Env) in which the trait was tested are displayed. AG: Algarrobo. AGHB4: *HaHB4*-introgressed line derived from Algarrobo.

The spike dry weight per m^2^ ranged from 55 to 191 g m^-2^ (AG in PE22RF and PE22IR2, respectively, Supplementary Table S6), while the fruiting efficiency varied from 76 to 249 grains g^-1^ (AGHB4 in PE22RF and PE18DS4, respectively, Supplementary Table S6). Both traits depended on the environment, but only the first one showed a trend (p=0.0787) to be affected by the environment × cultivar interaction when all data were considered (Supplementary Table S6). When the analyses were performed separately for environments where RY>0 or RY<0, a clear trend of the cultivar was detected for spike dry weight (p=0.0753 RY>0 and p=0.0797 RY<0), while no cultivar impact was computed on fruiting efficiency in any situation (Fig. 5C and D, Supplementary Table S6). The spike dry weight was 7.6% higher in AGHB4 than in AG when RY>0. In contrast, when RY<0, it was 12.8% lower in AGHB4 than in AG (Supplementary Table S6). The reduced spike dry weight of AGHB4 can explain the trend to lower GN and yield observed in environments with no WDE (Supplementary Tables S4 and S5). Despite the total biomass accumulated at anthesis seemed to be higher in AGHB4 than in AG when RY>0 (614 vs. 574 g m^-2^), and *vice versa* when RY<0 (863 vs. 946 g m^-2^), it was statistically affected only by the environment (Supplementary Table S6). The biomass partitioning to the spikes at anthesis was only affected by the environment exhibiting no trend between cultivars (Supplementary Table S6).

#### The AGHB4 used more water during the reproductive phase and showed higher water use efficiency than AG

The water used by the crop from emergence to maturity (WU_E-YP_) was only influenced by the environment, with no impact of the cultivar (Supplementary Table S7). It ranged from 262 (PE22RF) to 821 mm (PE22IR2), and it was ∼383 mm for both cultivars (average of all environments). The WUE_BT_ varied from 12.1 (AGHB4 in PE22RF) to 61.1 kg ha^-1^ mm^-1^ (AGHB4 in PE18SD3), while the WUE_YIELD_ ranged from 6 (PE22IR2 for AGHB4) to 28.8 kg ha^-1^ mm^-1^ (P18SD2 for AG). In contrast to WU_E-YP_, the WUE_BT_ and WUE_YIELD_ depended on both the environment and the cultivar, particularly when the RY>0 (Supplementary Table S7). In this condition, AGHB4 presented a 6.8 and 9.7% larger WUE_BT_ and WUE_YIELD_, respectively, than AG (Fig. 6A and B).

**Fig. 6.**
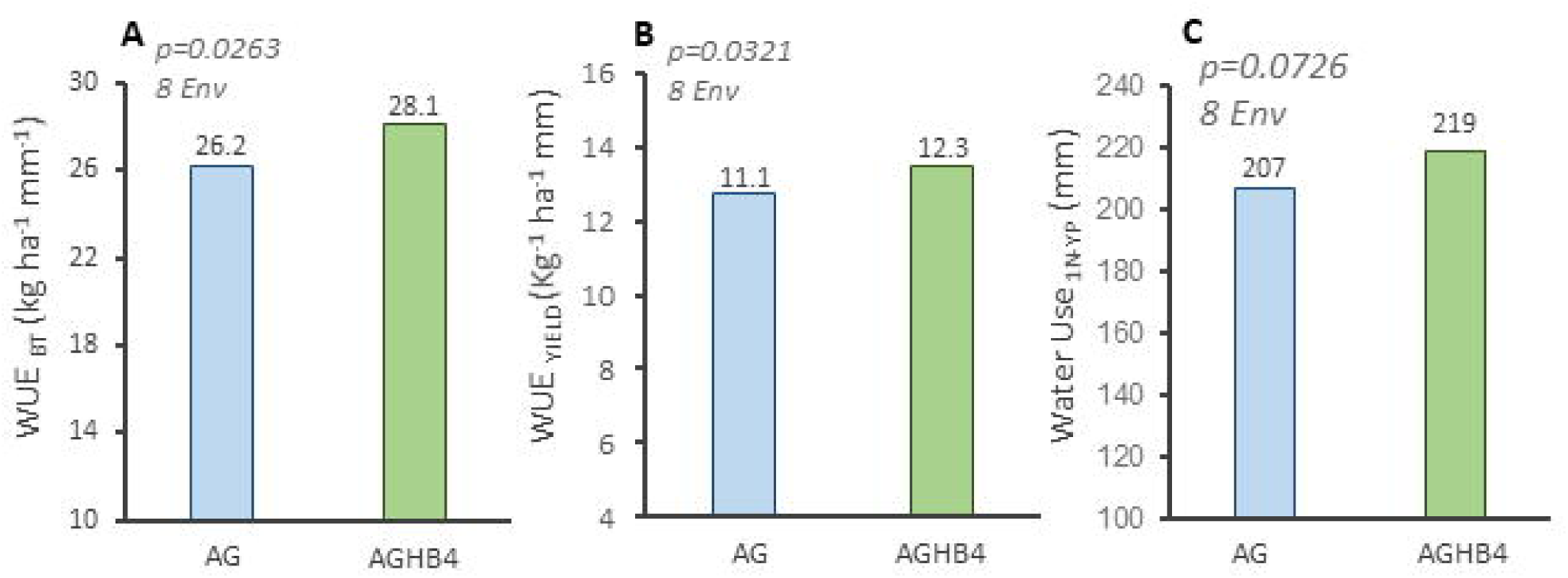
Water use (WU) and water use efficiency (WUE) in the environments where the relative yield (RY) of AGHB4 over AG was >0. (A) WUE for total biomass at maturity (WUE_BT_), (B) WUE for yield generation (WUE_YIELD_), and (C) water use during the reproductive phase from 1 node detectable (1N) to maturity (or yellow peduncle YP). The numbers above the bars indicate the exact value of the variable. In the upper left corner of each sub-figure, the p-value and the number of environments (Env) in which the trait was tested are displayed. AG: Algarrobo. AGHB4: *HaHB4*-introgressed line derived from Algarrobo.

Water use computed for the reproductive phase from first node detectable to maturity (WU_1N-YP_), when the impact of direct soil evaporation maybe reduced, depended not only on the environment but also on the cultivar, when RY>0. In this situation, AGHB4 showed a 5.5% greater WU_1N-YP_ than AG (Fig. 6 and Supplementary Table S7). It is important to note that both GN and yield showed a tight correlation with the WU_1N-YP_ (r=0.85 p<0.00001 and r=0.75 p<0.00001, for GN and yield, respectively).

As previously mentioned, 2022 was an extremely dry year in Pergamino, where we could set three water levels, rainfed (RF) and two irrigations (IR1 and IR2) due to the reduced rainfall during the crop cycle (Fig. 7A). In this environment, the canopy cover measured through the NDVI along the crop cycle tended to be greater in AGHB4 for all the water treatments, but particularly during pre-anthesis under the RF condition (Fig. 7B). The canopy temperature varied with the environment and the cultivar. As expected, the IR2 and IR1 showed lower canopy temperatures than the RF (Fig. 7C). AGHB4 presented, in general, a cooler canopy than AG, being the Canopy Temperature Difference (CTD) between cultivars greater under the RF than the IR1 and IR2 conditions (Fig. 7C and 7D). The particularly dry condition of the 2022 year allowed us to detect a trend to different soil water uptake in the soil profile between the cultivars (Supplementary Fig. S5). On the one hand, AGHB4 tended to extract more water from the 1 to 2 m soil depth under RF conditions, visualized as a slight trend at 1 Node but particularly evident at maturity (Supplementary Fig. S5A and S5B). On the other hand, no difference was observed at maturity between cultivars under full irrigation (IR2), but AGHB4 tended to extract more water than AG from the first 1 m soil depth when there was a high supply in this soil layer (Supplementary Fig. S5C and S5D). Collectively, the results about CTD and soil water depletion support the findings referred to water use.

**Fig. 7.**
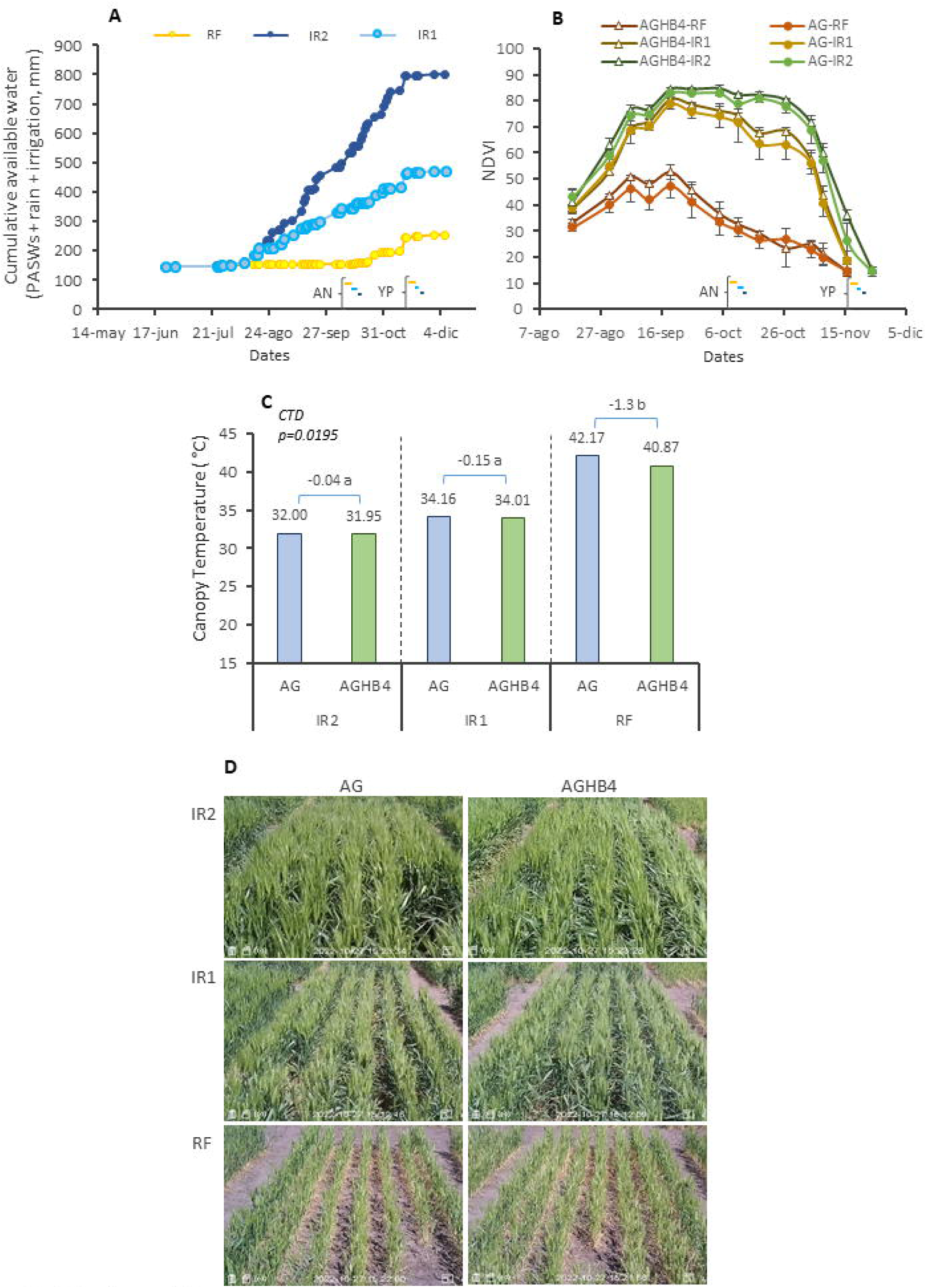
Pergamino 2022 environment description and its impact on canopy cover and temperature. (A) Cumulative available water for crop growth considering the plant available soil water at sowing up to 2m depth (PASWs), the rain and irrigation for the rainfed condition (RF), and the two irrigation levels (IR1: 220mm -IR2 548mm). (B) Normalized vegetation index (NDVI) during crop cycle for the three water conditions (RF, IR1 and IR2) and both cultivars. The coloured lines at the bottom of (A) and (B) indicate the range in dates of anthesis (AN) and yellow peduncle (YP) under each water condition, considering both cultivars. (C) Canopy temperature for each combination of water treatment and cultivars. The canopy temperature difference (CTD) between cultivars for each water condition is presented. Different letters indicate statistical differences (Tukey α=0.05). (D) Illustrative pictures of the plots of AG and AGHB4 under RF, IR1 and IR2, ∼10 days post-anthesis, on 27 October. AG: Algarrobo. AGHB4: *HaHB4*-introgressed line derived from Algarrobo.

## Discussion

### Yield advantage of HaHB4 on a modern cultivar adapted to the growing conditions

The average yield advantage of expressing *HaHB4* in Algarrobo was 3.3% in the greenhouse and 8.1% in the field, the latter being 2.1% higher than the benefit observed in Cadenza-HB4 (6%) (González et al., 2019). When water deficit during the reproductive phase was present, the gain rose to 15% in the greenhouse, and 13% in the field (computing WDE during the months SON or OND according to the location). These grains align with the 16% yield benefit in Cadenza-HB4 in dry environments (González et al., 2019), confirming no overestimation of *HaHB4* benefit due to Cadenza’s long cycle. The absence of a significant background effect on the *HaHB4* yield advantage was corroborated by these findings, which is crucial for the widespread adoption of this technology. Similar to González et al. (2019), few environments (4 out of 27) yielded no advantage of *HaHB4* introgression, particularly in non-water-deficit environments, where a non-significant trend to higher AG yield was absorbed. This was clear in well-watered greenhouse conditions and when considering the plant available soil water at the beginning of the reproductive period in the water balance (see Fig. 3C). These results underscore the significant role of the sunflower transcription factor *HaHB4* under water deficit, while also highlighting its lack of impact in non-water-deficit environments.

In this study, we fitted a response curve for the RY of AGHB4 compared to AG in relation to the water deficit during the reproductive phase (see Fig. 3B and C). The introgression of *HaHB4* improved RY by 0.06 to 0.08 % per mm of water deficit. This analysis represents a preliminary quantification of the benefit of *HaHB4*-wheat crop in environments with varying water deficit levels. For instance, for the 2000 to 2020 climatic series, the 50% probability of WDE during the reproductive phase ranged from −155 mm in Roldán to −264 mm in Bordenave (Supplementary Fig. S6). Considering a benefit of 0.06% per mm, the yield gain from *HaHB4* would range from 9.3% to 15.8%. These calculations apply when the crop enters the reproductive phase with a dry soil. In wetter years, the plant available soil water may mitigate the deficit, but the introduction of *HaHB4* would still provide benefits (Supplementary Fig. S6) unless the WDE during the reproductive phase reaches zero (Fig. 3C). Crop simulation models will be used in an upcoming paper to more accurately assess the benefits of the *HaHB4* introgression in various wheat production areas. We hypothesize that the unique regression presented in this paper actually represents a family of curves that vary with soil types (and water retention, which would modify the x-axis) as well as with heat stress temperature and atmospheric vapor pressure deficit (potentially resulting in different yield advantages per mm of water use as they influence the WUE, Hatfield and Dold, 2019).

In 2022, Pergamino experienced an extremely dry and warm season, resulting in a lower yield advantage for *HaHB4* than expected, particularly under high water deficit (see underlined dots in Fig. 3B and C). In our previous work (González et al., 2020), dry environments with warm temperatures during the reproductive phase (>20 °C mean temperature) raised yield improvement to 20%. A similar response was reported in Williams 82 soybean expressing the *HaHB4* gene, increasing benefit to 11% when dry and warm environments were considered (González et al., 2020). In the present paper, all environments explored mean daily temperatures > 20 °C, and daily maximum temperatures > 30 °C during the reproductive phase (Supplementary Table S1). To asses the impact of these temperatures, temperature indexes were calculated summing up daily mean temperatures > 20 °C or maximum mean temperatures > 30 °C during the reproductive phase (SON/OND, see Supplementary Table S1). The relative yield of AGHB4 compared to AG exhibited an optimal response curve to both thermal indices; however, the ∑Tmax > 30 °C index, representing a heat stress, accounted for greater variability (Fig. 8A). When the residuals from the regression of the RY of AGHB4 compared to AG against the WDE (Fig. 3B) were related to this heat stress index, a similar optimal response curve emerged (Fig. 8B). This suggests that *HaHB4* was responsive not only to water deficit but also to moderate heat stress, which may account for part of the variability in its response. The increased water use, leading to a reduction in canopy temperature when *HaHB4* is introgressed (Fig. 6 and 7), may be partially mediating this response. These novel findings encourage further experiments given the more frequent occurrence of combined drought and high temperature events in recent years across many regions around the world (Mukherjee and Mishra, 2021).

**Fig. 8.**
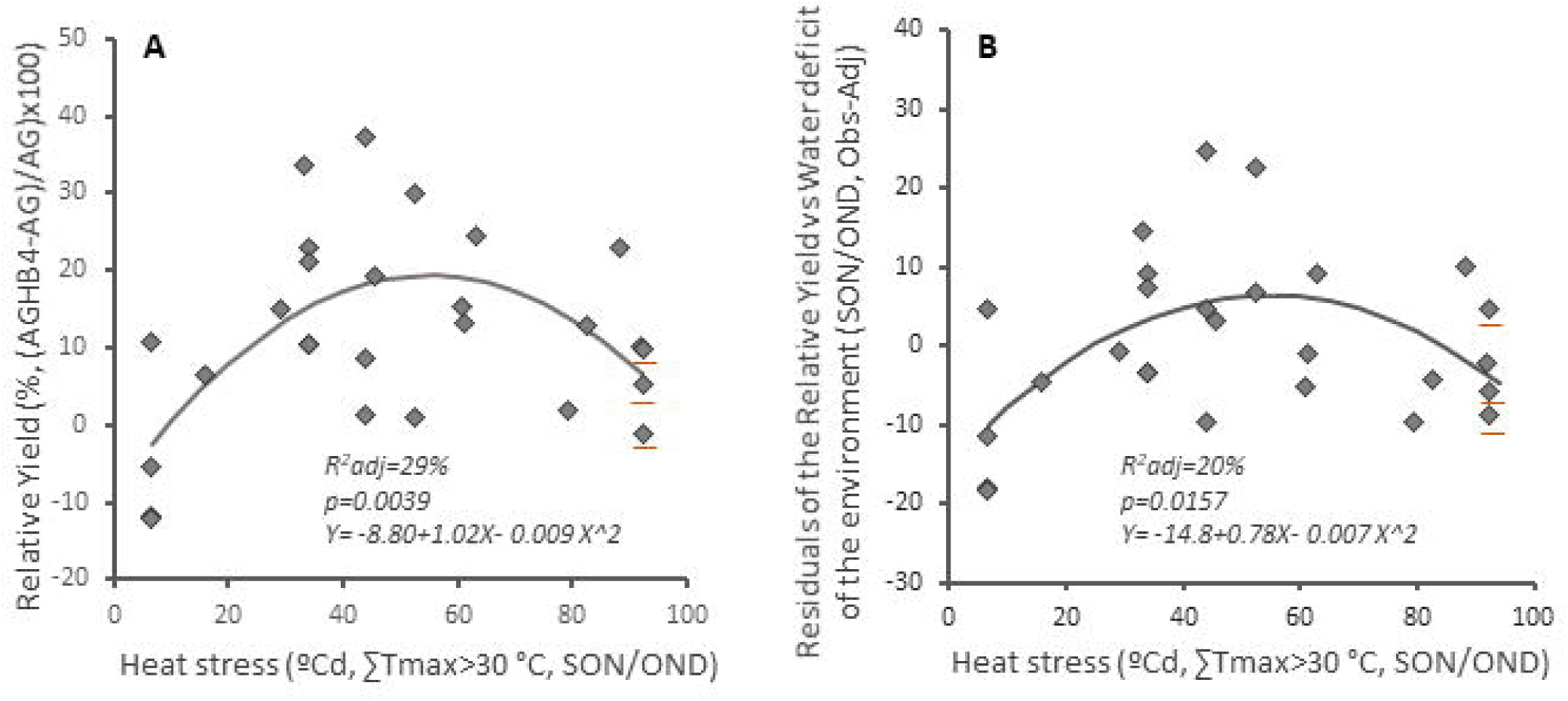
Response to heat stress of (A) the relative yield of AGHB4 over AG and (B) the residuals of the regression between the relative yield of AGHB4 over AG and the water deficit of the environment during the months when the reproductive phase takes place (September, October and November -SON- for northern locations and October, November and December – OND for southern locations). Obs-Adj: Observed - Adjusted. ∑ Tmax>30 °C: daily sum of maximum mean temperature >30 °C during SON/OND. The three underlined dots represent the response for PE2022 rainfed and the two irrigation levels. AG: Algarrobo. AGHB4: *HaHB4*-introgressed line derived from Algarrobo.

### Physiological process underpinning the yield advantage of the HaHB4 introgression under water deficit

As anticipated in González et al. (2019), no relevant impact on crop phenology was observed. The yield benefit from *HaHB4* was mainly associated with the number of grains per m^2^, consistent with Cadenza’s response (González et al., 2019), and reinforcing the well-established importance of this component in yield determination (Fischer, 1985; Miralles and Slafer, 2007; Curin et al., 2021). The number of spikes per m^2^ at anthesis, resulting from enhanced tiller survival, were the primary driver of improved grains per m^2^ (Fig. 9). This is a novel finding as previous work with Cadenza-HB4 detected greater number of spikes per plant in one environment (González et al., 2019). Heavier spikes per m^2^ at anthesis accompanied the enhanced grains per m^2^ in AGHB4 (Fig. 9), contrasting with spike-level results in Cadenza-HB4 leading to more fertile florets per spike (González et al. 2019). While no significant changes in biomass allocation to reproductive spikes at anthesis were noted, a slight but statistically significant increase in the HI was observed in AGHB4. Previous work with Cadenza-HB4 showed variable response in GW, whereas in the present study, heavier grains were detected in AGHB4 (Fig. 9). The difference between Algarrobo and Cadenza background in the relative importance of yield components can be attributed to the environments experienced. Each numerical component exhibits different plasticity, being spikes per m^2^ > grain per spikes > grain weight (Sadras and Slafer, 2012). Cadenza’s longer cycle may have exposed the reproductive phase to poor environments, limiting tillers plasticity, and consequently, the response of spikes per m^2^. However, the increased fertile florets per plant in Cadenza-HB4 (González et al., 2019) aligns with the enhanced tiller survival and spike growth observed in AGHB4, as both depend on the crop growth from first detectable node to anthesis (Fischer, 1985; Slafer et al., 2005). The greater crop growth rate and BT achieved in Cadenza-HB4 (González et al., 2019, 2020) was also observed in AGHB4, reinforcing the notion that the primary effect of *HaHB4* is to sustain crop growth under water deficit. Agreeing with these results, all the vegetation indices indicated a better crop condition of AGHB4 compared to AG under water deficit (see Fig. 2), while few of them differentiated both cultivars in well-irrigated conditions. In soybean, the benefits from *HaHB4* were also associated with improved GN due to greater crop growth and biomass production (Ribichich et al., 2020).

**Fig. 9.**
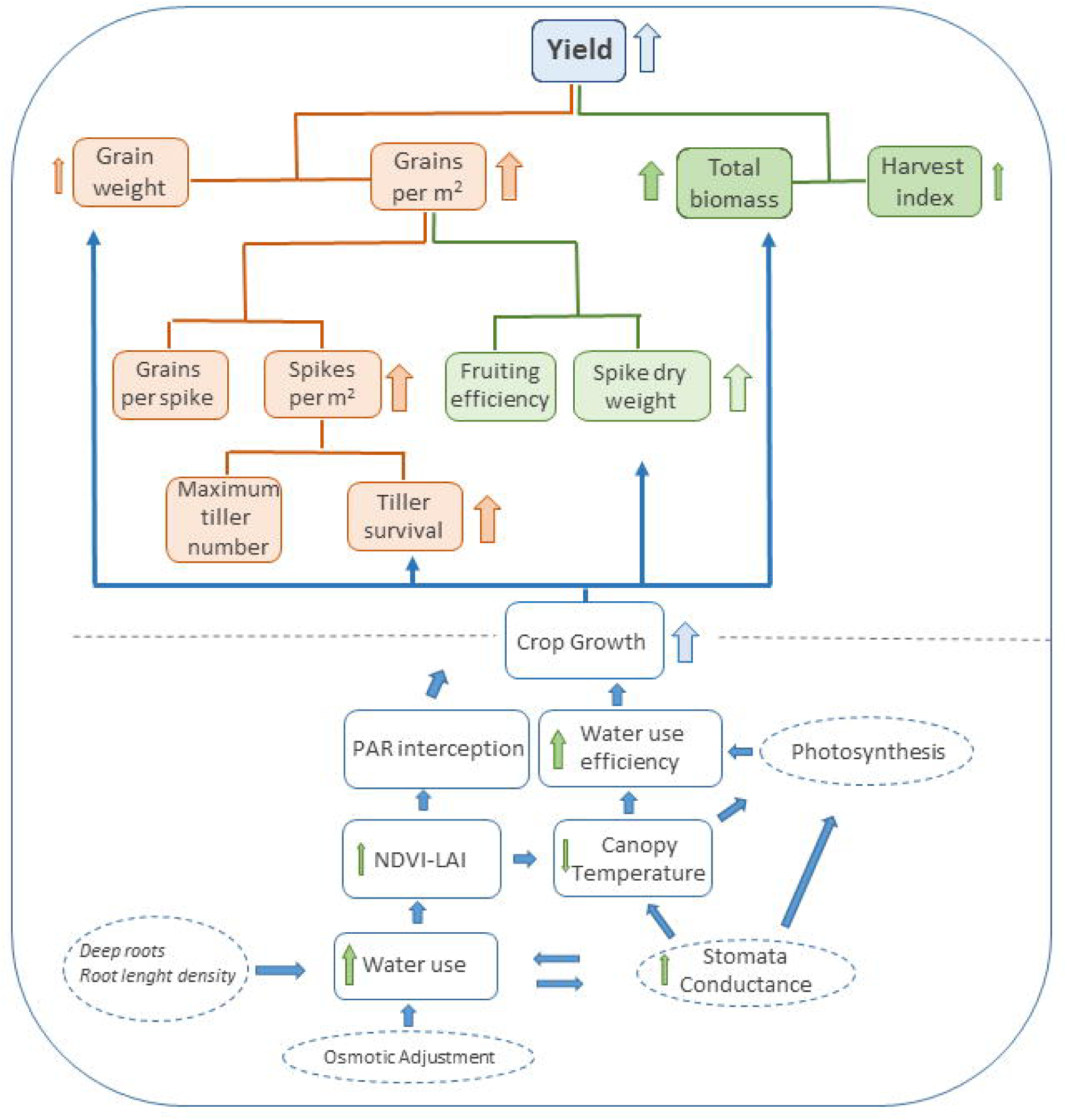
Schematic representation of possible processes determining the yield benefit under moderate/severe water deficit and moderate heat stress of *HaHB4*-introgression. The size of the arrows next to the traits represent the level of response observed. NDVI: Normalized Vegetation Index. LAI: leaf area index.

The WU is a key trait determining crop growth (Passioura, 1977, 1996) under water deficit. WU encompasses both the direct evaporation from the soil and the transpiration by the crop. As leaf area index increases, the relative weight of transpiration in defining the WU increases (Ritchie and Burnett, 1971). Therefore, WU during the reproductive period (WU_1N-YP_) more accurately reflects crop transpiration than WU throughout the entire crop cycle (WU_E-YP_). Under water deficit, WU_1N-YP_ was greater for AGHB4, with 21.9% and 5.5% increases in greenhouse and field conditions, respectively. In the case of soybean, the *HaHB4* line tested not only used more water under deficit (+17.3%), but also under irrigation (+27.2%) due to its larger hypocotyl diameter and xylem area, traits linked to enhanced water conductivity (Richards and Passioura, 1981). In contrast, the WU_1N-YP_ in AGHB4 was similar or lower than the control cultivar under well-watered conditions, suggesting a different mechanism. WU under water deficit would be affected by deep roots (Lopes and Reynolds, 2010), high root length density (Loomis and Connor, 1992) or osmotic adjustment which would help to absorb water from drying soils with low water potential (Tardieu, 2005; Lambers et al., 2008; Blum, 2017) (Fig. 9). Although a trend toward deeper soil water absorption was observed in AGHB4 during the driest year in Pergamino (see Supplementary Fig. 5), differences in water absorption were also detected in the 1 m depth greenhouse containers. The potential for high root length density or osmotic adjustment in the AGHB4 should not be overlooked. Further critical experiments are needed to definitively elucidate the mechanisms underlying the increased water use observed in the *HaHB4*-introgressed wheat.

The canopy cover (and interception of photosynthetic active radiation), as assessed by NDVI, was greater in AGHB4, particularly under water deficit in both greenhouse (Fig. 2C) and field conditions (Fig. 7B). Additionally, the leaf area index under water deficit in the greenhouse tended to be higher in AGHB4 (Supplementary Table 3). The maintenance of the leaf area, which mainly relies on the expansive growth (Tardieu et al., 2011), may have been supported by the higher WU_1N-YP_ observed in AGHB4 (Fig. 9). The reduced canopy temperature under water deficit in AGHB4 (Fig. 7C) aligned with the increased water use and the trend towards greater stomatal conductance) (Fig. 9). In soybean, enhanced radiation interception during the reproductive phase, also evidenced as prolonged stay green, and increased leaf photosynthesis conferred by *HaHB4* contributed to the biomass increment (Ribichich et al., 2020). In line with these results, in sunflower and Arabidopsis, *HaHB4* acted repressing specific targets of the ethylene perception which was translated into a delay to entry senescence under water deficit (Manavella *et al.,* 2006). Conversely, no significant delay in senescence or notable impact on leaf photosynthesis was detected in AGHB4. A possible explanation may be the existence of divergent pathways involving ethylene response between dicotyledonous and monocotyledonous plants, as occurs in plants growing in darkness (Satler and Kende, 1985; Ishizawa and Esashi, 1988; Lee and Lin, 1996; Yang *et al.,* 2015). The *HaHB4* strategy to cope with water deficit appears to be growth maintenance via increasing water uptake, a valid approach for climatic scenarios with mild water deficits (Tardieu, 2005).

The WUE_BT_ was higher in the AGHB4 under water deficit in filed conditions, consistent with findings in the HB4-soybean study (Ribichich *et al.,* 2020). This was noted despite the higher water use in both species when *HaHB4* was introgressed. An improved water use efficiency at the leaf level, or transpiration efficiency, calculated as the quotient of maximum photosynthesis to stomatal conductance (Hatfield and Dold, 2019), may have contributed to this result. HB4-Soybean exhibited a higher photosynthesis rate during grain filling, likely linked to increased stomatal conductance and stay green processes (Ribichichi et al., 2020). Nevertheless, a direct impact of *HaHB4* on photosynthesis, should also be considered. Photosynthesis-related gene expression was altered by the *HaHB4* in Arabidopsis; the transcription of photosynthesis genes in darkness was inhibited leading to energy saving without a decay in the photosynthesis rate (Manavella et al., 2008). Surprisingly, the enhanced WUE_BT_ in AGHB4 was detected only under field conditions with no impact in the greenhouse. The temperature in the greenhouse was set to 25 °C, which is the optimum temperature for wheat leaf photosynthesis (Kobza and Edwards, 1987). Under these conditions no positive trend to improve photosynthesis by *HaHB4* was observed. Though leaf photosynthesis was not measured in current research in the field, we hypothesize that the reduced canopy temperature in the AGHB4, resulting from higher water use under water deficit, may have contributed to lowering leaf temperature, thereby mitigating the possible reduction in photosynthesis rate due to temperatures higher than 25 °C. This mechanism may also partially explain the moderate heat stress tolerance observed in AGHB4. The improvement in WUE_YIELD_ observed in AGHB4 was also reported for soybean expressing *HaHB4* gene (Ribichich et al., 2020) and is in line with the higher efficiency to produce yield per mm of rainfall of Cadenza-HB4 (González et al., 2019). The enhancement of HI, triggered mainly by the GN and to a lesser extent by GW, accounted for increased WUE_YIELD_ in AGHB4. The increase in WUE through breeding, which has been identified as a strategy to mitigate the impact of more severe drought conditions (Tardieu, 2005), would expand the range of environments where *HaHB4* can provide yield advantages.

## Conclusions

Under water deficit during the reproductive phase, the introgression of *HaHB4* into Algarrobo showed an average yield advantage of 15% in the greenhouse and 13% in the field. Since similar results were observed in Cadenza’s background, we conclude that the benefits of *HaHB4* were not overestimated in our previous tests due to Cadenza’s long growth cycle when compared to well-adapted cultivars. *HaHB4* improved the relative yield by 0.06 to 0.08 % per mm of water deficit compared to its control counterpart. This absence of a significant background effect on the HaHB4 yield benefit is crucial for the widespread adoption of this technology. *HaHB4* was also responsive to temperatures above 30 °C, showing a yield advantage under moderate heat stress. The yield advantage was a consequence of the sustained growth under water deficit associated to higher water use during the reproductive period and improved water use efficiency to produce biomass and yield. *HaHB4* would improve the water-limited yield in moderate to severe drought environments, enhancing wheat yield stability and reducing the gap with the potential yield. This may have a great impact in productivity in rainfed cropping systems, like most of the wheat producing areas around the world.

## Supplementary Data

The following Supplementary data are available at *JXB online*

**Fig. S1.** Comparison between Cadenza and Algarrobo.

**Fig. S2.** Schematic representation of plant material generation and locations where experiments were carried out.

**Fig. S3**. Greenhouse containers. Pictures and evolution of total water (mm) up to 1m depth.

**Fig. S4**. Phenology of Algarrobo (AG) and its *HaHB4*-introgressed line (AGHB4) at different sowing dates.

**Fig. S5**. Water soil profiles in Pergamino 2022.

**Fig. S6**. Accumulated probability of water deficit of the environment and relative yield advantage of AGHB4 over AG.

**Table S1**. Description of environments explored, sowing and anthesis dates for each environment.

**Table S2**. List of selected Vegetation Indices.

**Table S3.** Response to well-watered (WW) and water-deficit (WD) regimes of yield and its numerical components and crop physiological determinants.

**Table S4.** Yield of Algarrobo (AG) and its *HaHB4*-introgressed line (AGHB4) in the different environments.

**Table S5.** Numerical components of yield for Algarrobo (AG) and its *HaHB4*- introgressed line (AGHB4).

**Table S6.** Crop physiological determinants of yield in Algarrobo (AG) and its *HaHB4*- introgressed line (AGHB4).

**Table S7.** Water use and water use efficiency in Algarrobo (AG) and its *HaHB4*- introgressed line (AGHB4).

## Acknowledgments

We would like to thank Luis Blanco and Octavio Ghio Trebino for field technical assistance. We also acknowledge Marcos Pareta, Matías Martin, Matías Cardoso Perassi and Joaquín Montiel Moyano who prepare their undergraduate thesis within the development of some of the experiments of the present study.

## Author contribution

FA, MDZ, FC and FG performed the experiments, MP and EBM took, analyzed and wrote the section about hyperspectral data and vegetation indices, FG analyzed the rest of the agronomic and physiological data, with assistance of FC, JM, and RC helped with the molecular aspects of the introduction and discussion, FG designed the study, FG and RC conceived the ms, FG wrote de ms, MP, MO, RC and FA help with the general writing. All authors revised and approved the ms. FA and FC contributed similarly to this work.

## Conflict of interest

The authors declare no conflict of interest. FA is current employee of Bioceres Crop Solutions.

## Funding

This work was funded by Agencia Nacional de Promoción de la Investigación, el Desarrollo Tecnológico y la Innovación (Agencia I+D+i, PICT2015-2671) competitive grant.

## Data Availability

Most data are available in the Supplementary Tables in *JXB online*. Further information can be provided upon request.

## Abbreviations

AG: Algarrobo cultivar
AGHB4: *HaHB4*-introgressed line derived from Algarrobo cultivar
BT: total biomass at maturity
GW: individual grain weight
GN: grain number per m^2^
HI: harvest index
IR: drip irrigation
RF: rainfed
RY: relative yield
WD: water deficit
WDE: water deficit of the environment
WU: water use
WUE: water use efficiency
WUE_BT_: water use efficiency to produce total biomass at maturity
WUE_YIELD_: water use efficiency to produce yield
WW: well-watered

## Notes

### Competing Interest Statement

The authors have declared no competing interest.

